## Supplementary figures and images for "Cerebellar nuclei cells produce distinct pathogenic spike signatures in mouse models of ataxia, dystonia, and tremor"

### Figure 1 - Supplementary Figure 1

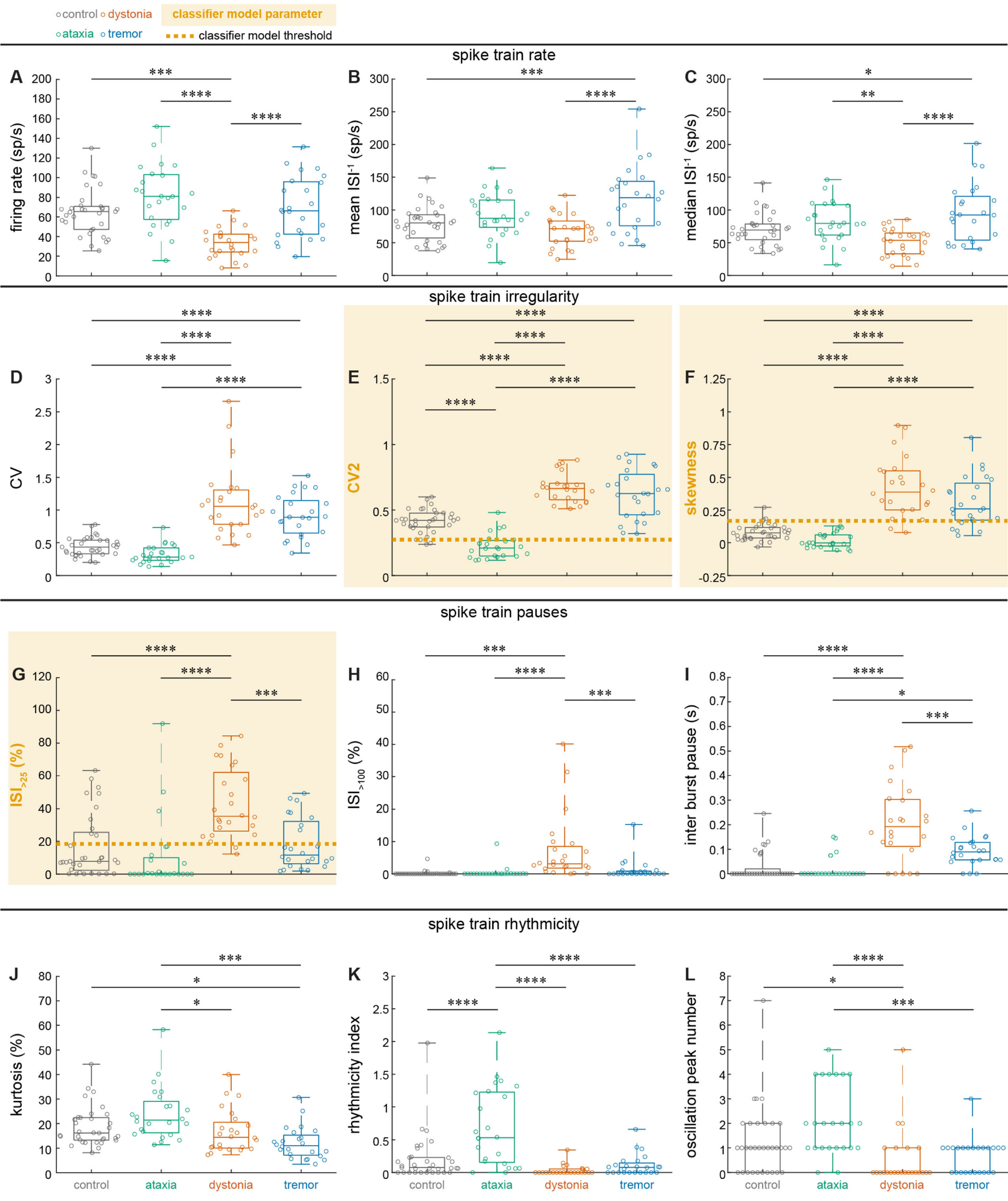

### Figure 1 - Supplementary Figure 2

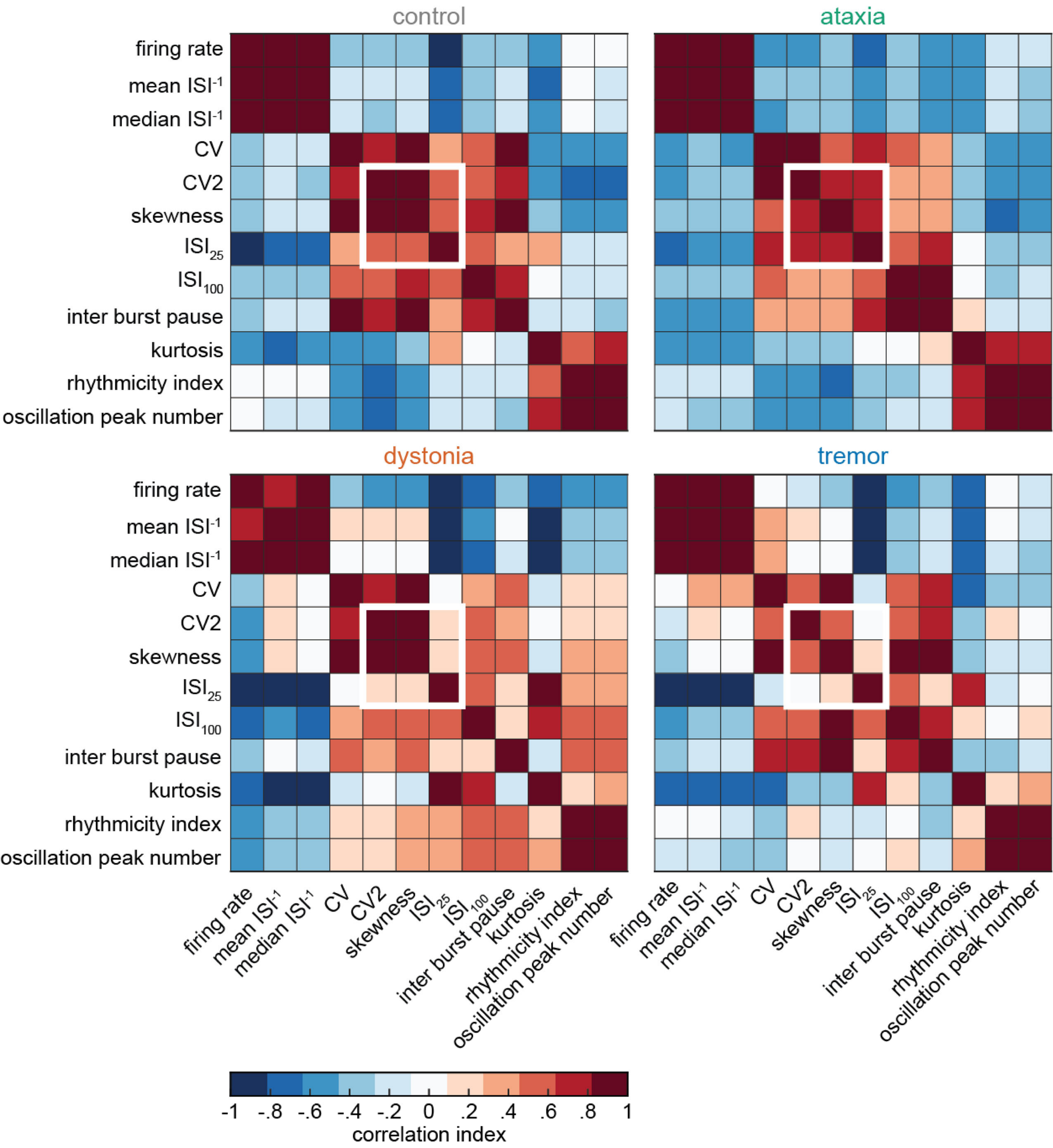

### Figure 1 - Supplementary Figure 3

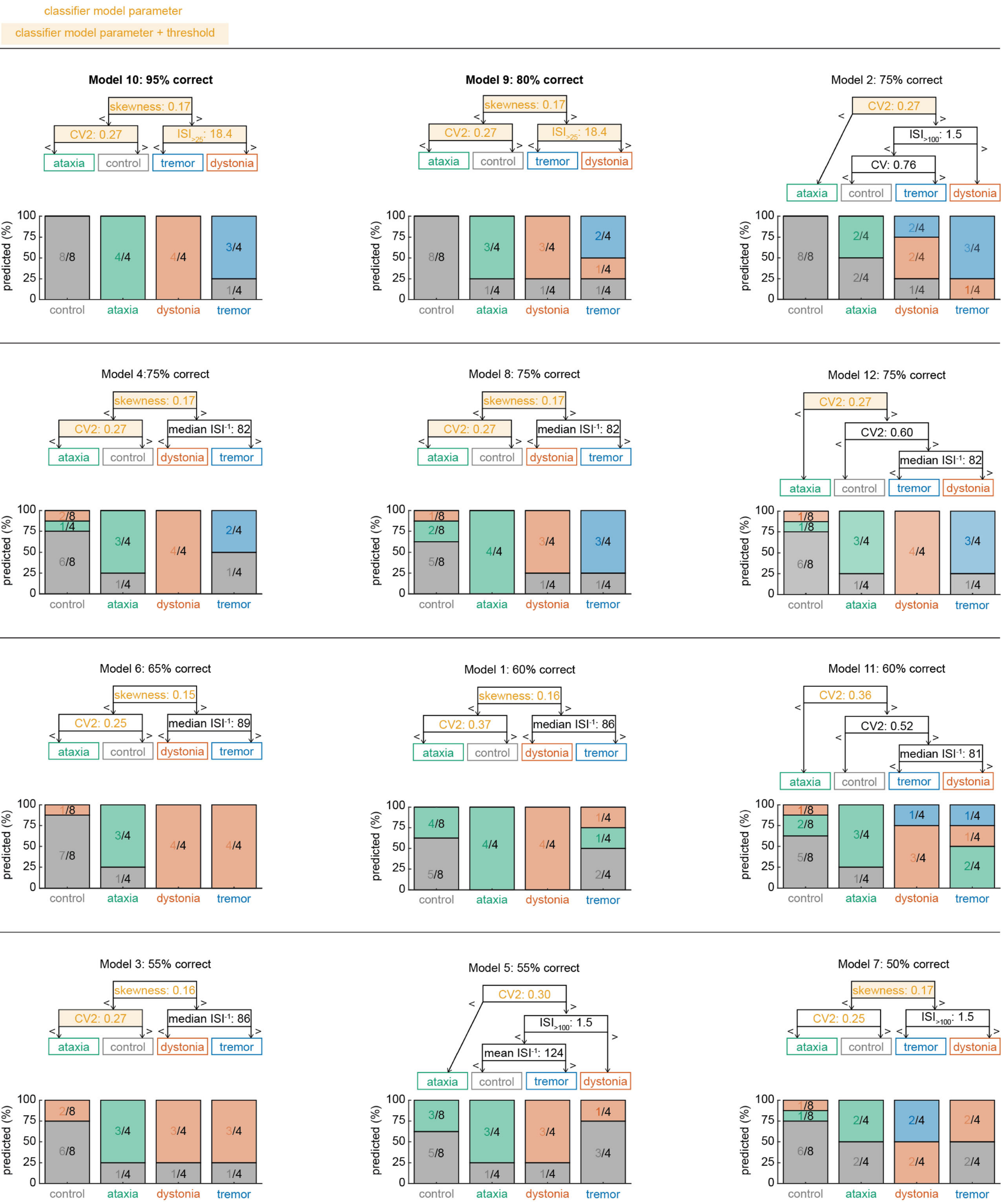

### Figure 5 - Supplementary Figure 1

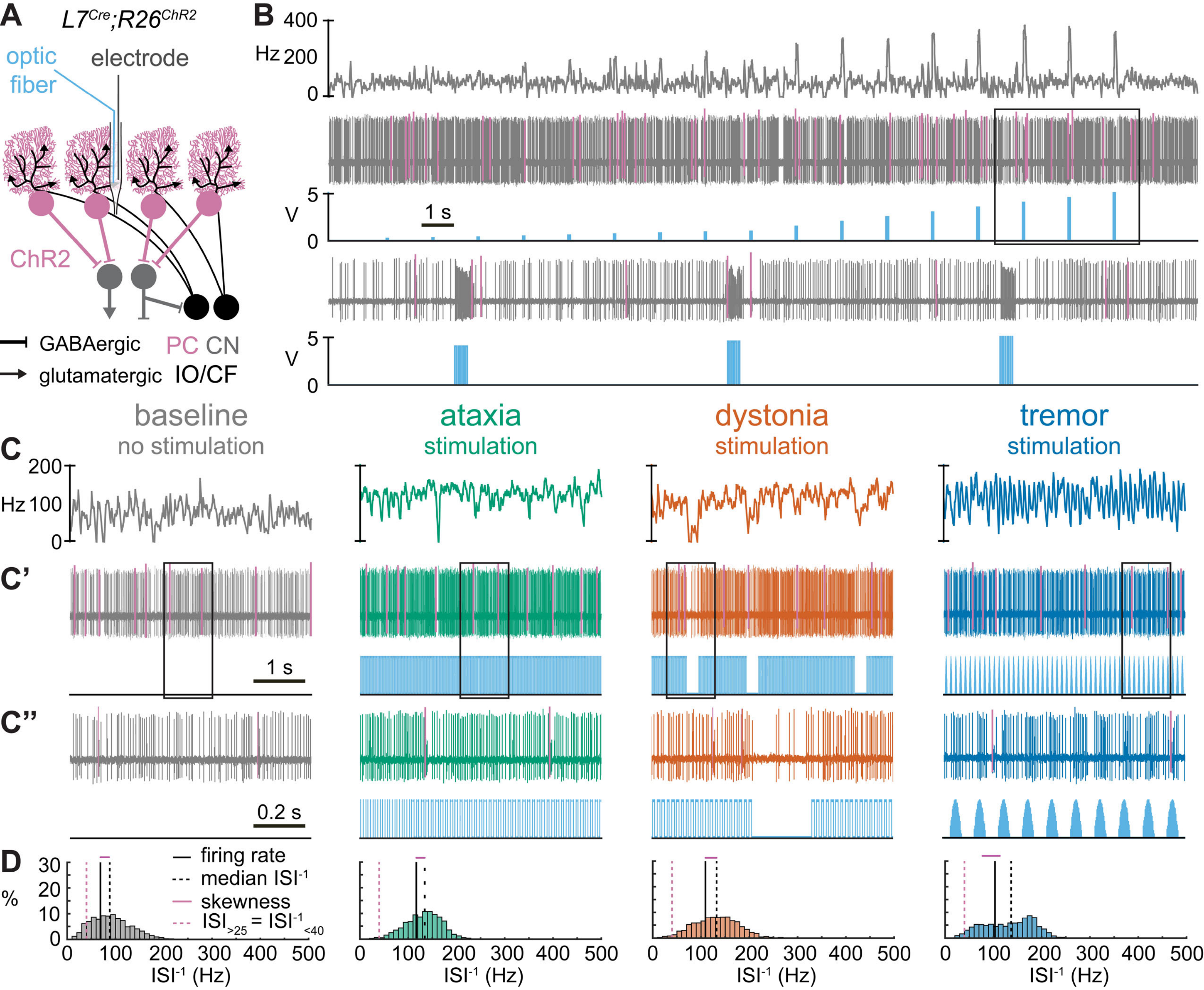

### Figure 5 - Supplementary Figure 2

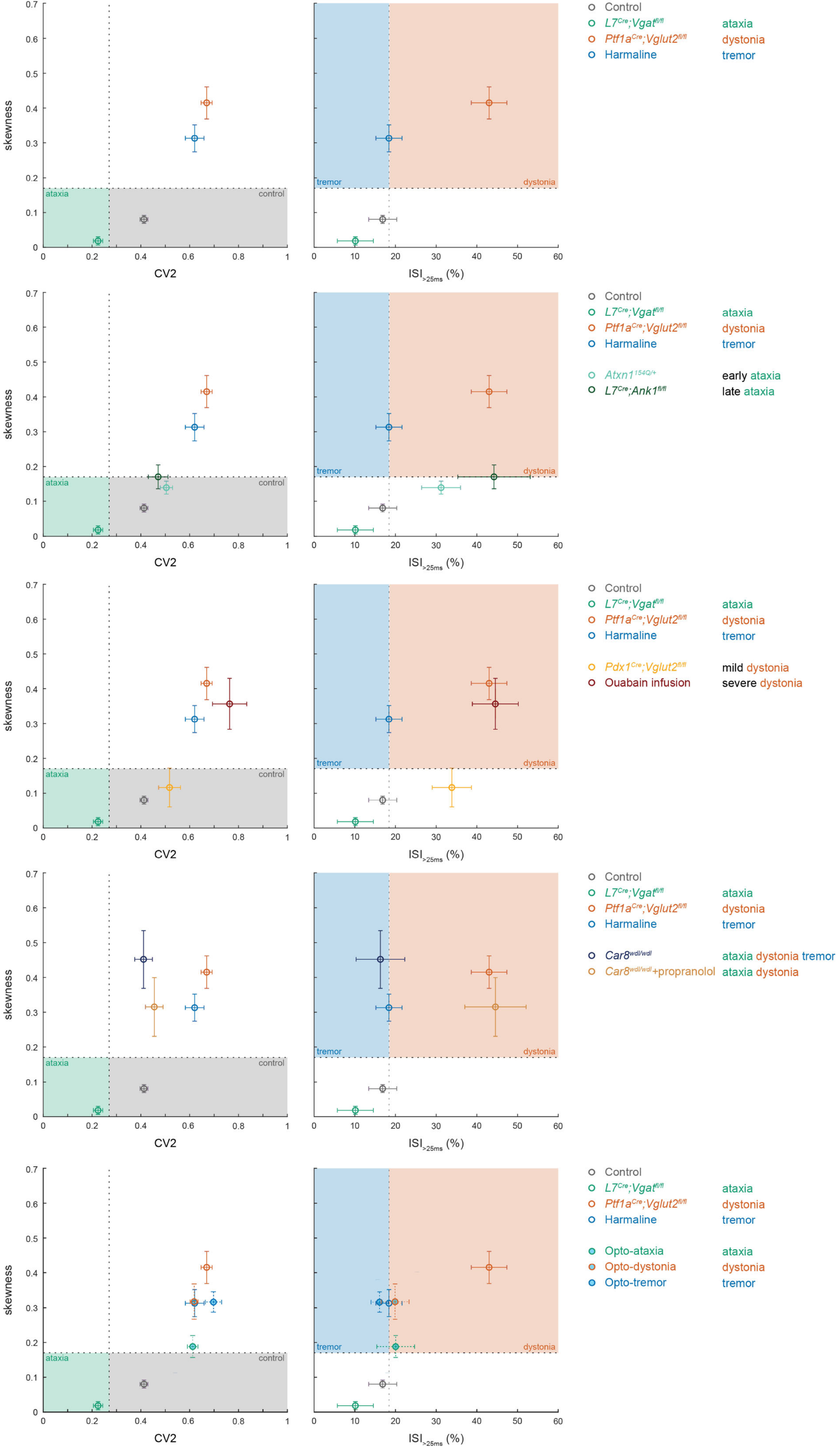
